# An evaluation of transcriptome-based exon capture for frog phylogenomics across multiple scales of divergence (Class: Amphibia, Order: Anura)

**DOI:** 10.1101/031468

**Authors:** Daniel M. Portik, Lydia L. Smith, Ke Bi

## Abstract

Custom sequence capture experiments are becoming an efficient approach for gathering large sets of orthologous markers with targeted levels of informativeness in non-model organisms. Transcriptome-based exon capture utilizes transcript sequences to design capture probes, often with the aid of a reference genome to identify intron-exon boundaries and exclude shorter exons (< 200 bp). Here, we test an alternative approach that directly uses transcript sequences for probe design, which are often composed of multiple exons of varying lengths. Based on a selection of 1,260 orthologous transcripts, we conducted sequence captures across multiple phylogenetic scales for frogs, including species up to ~100 million years divergent from the focal group. After several conservative filtering steps, we recovered a large phylogenomic data set consisting of sequence alignments for 1,047 of the 1,260 transcriptome-based loci (~630,000 bp) and a large quantity of highly variable regions flanking the exons in transcripts (~70,000 bp). We recovered high numbers of both shorter (< 100 bp) and longer exons (> 200 bp), with no major reduction in coverage towards the ends of exons. We observed significant differences in the performance of blocking oligos for target enrichment and non-target depletion during captures, and observed differences in PCR duplication rates that can be attributed to the number of individuals pooled for capture reactions. We explicitly tested the effects of phylogenetic distance on capture sensitivity, specificity, and missing data, and provide a baseline estimate of expectations for these metrics based on nuclear pairwise differences among samples. We provide recommendations for transcriptome-based exon capture design based on our results, and describe multiple pipelines for data assembly and analysis.

## Introduction

Using high throughput sequencing, there are now a variety of approaches available to generate large molecular data sets for the purpose of addressing population genetics or phylogenetics questions. A majority of these approaches fall in the category of reduced representation sequencing, in which orthologous sets of markers from a subset of the genome are obtained across taxa or individuals. A commonly used approach is RAD-seq, which targets anonymous loci adjacent to restriction enzyme sites (Miller et al. 2007), though the probability of obtaining orthologous sets of loci decreases as phylogenetic distance between samples increases (Rubin et al. 2012; Arnold et al. 2013). Other approaches include more targeted selection of loci using DNA or RNA probes, such as ultra-conserved element (UCE) sequencing (Faircloth et al. 2012) and anchored hybrid enrichment (Lemmon et al. 2012). Both approaches rely on short, highly conserved genomic regions for probe design and the subsequent capture of these targets for libraries with large insert sizes containing stretches of flanking sequences. This allows the use of the same set of markers across distantly related taxa, but the function of these loci is generally unknown, and the levels of variation in flanking regions are not predictable. Other targeted sequence capture approaches allow more control over the level of variation of orthologous markers, including sequence capture using PCR-generated probes (SCPP) (Peñalba et al. 2014), and transcriptome-based exon capture (Bi et al. 2012). The latter approach uses transcriptome sequencing to identify protein-coding exons across populations or species, and is particularly useful for organisms for which no other genomic resources are readily available.

An important step before selecting markers derived from transcriptome sequences involves the identification of intron-exon boundaries to select longer exons, which requires the use of reference genomes (Bi et al. 2012; Bragg et al. 2015). Longer exons are preferred because they exceed the length of capture probes, allowing tiling, and for a given evolutionary rate they should have more informative sites compared to shorter exons. The transcriptome sequences recovered are typically composed of multiple exons, often short in length, making probe design challenging. The intron-exon identification step can be exceedingly difficult if the reference genome is too distantly related, and the direct use of transcriptome sequences for probe design is an alternative that has not been explored. Although this alternative approach ignores the potential presence of intron-exon boundaries, it offers an opportunity to capture exons of a variety of lengths along with their associated non-coding flanking regions. The length of probes, level of divergence between probes and targets, length distribution of genomic library fragments, and the number of and lengths of exons in the transcript sequences are all important factors that could determine the success of this alternative approach.

There are several major challenges for designing a custom sequence capture experiment for a non-model organism, particularly if the experiment involves species with relatively large genome sizes, spans multiple phylogenetic scales, and involves the *de novo* generation of genetic resources for probe design. In addition, wet-lab-specific decisions have the potential to significantly influence the outcome of sequence captures, including the number of genomic libraries to pool per capture reaction and the choice of genomic library blocking oligos. Few studies have focused on the exploration of these topics across a single experiment, yet the availability of baseline information can help inform these decisions and improve the success of sequence capture.

Across terrestrial vertebrates, amphibians exhibit the largest genome sizes. The average genome size of frogs is 5.0 gigabases (Gb) (max = 13.1 Gb, n = 497), whereas the salamander genome averages 34.5 Gb (max = 117.9 Gb, n = 426) (Gregory 2015). These genome sizes are larger than those of birds (1.3 Gb, n = 896), mammals (3.1 Gb, n = 777), and squamates (2.1 Gb, n = 344) (Gregory 2015), and the performance of targeted exon capture for amphibians remains largely unexplored (but see Hedtke et al. 2013; McCartney-Melstad et al. 2015). Here, we examine the performance of transcriptome-based exon capture for frogs across multiple phylogenetic scales. The main focal group is the African frog family Hyperoliidae, consisting of 13 genera and 254 samples, which have an average genome size of 4.6 Gb (n = 11) (Gregory 2025). Our sampling also includes species from the sister family Arthroleptidae (7 genera, 7 samples), and a single representative from three more distantly related families (Brevicipitidae, Hemisotidae, Microhylidae). Pairwise comparisons within Hyperoliidae do not exceed 10% nuclear divergence, and the hyperoliid genera shared a common ancestor approximately 56 million years ago (Portik & Blackburn, in prep). The family Hyperoliidae shares a common ancestor with Arthroleptidae approximately 77 Ma, with Hemisotidae and Brevicipitidae approximately 93 Ma, and with Microhylidae approximately 103 Ma, and uncorrected pairwise nuclear differences between hyperoliids and the outgroups approaches 20%.

We describe our methodological approach for generating and mining transcriptome resources, the selection of orthologous markers and probe design, choice of blocking oligos in capture reactions, the pipeline for assembling and processing capture sequence data, and the overall results of our exon capture experiment. Given the tremendous level of divergence between our focal group and available frog reference genomes (*Xenopus laevis* and *X. tropicalis,* minimum 150 million years divergent), we did not attempt to identify intron-exon boundaries to select longer exons. Rather, we use transcriptome sequences directly for probe design. We evaluate our results given this approach, including characterizing the number of exons in transcript sequences, the lengths of these exons, our ability to recover exons and their flanking regions, and the effects of exon length on sequencing depth. The level of variation within the family Hyperoliidae and the inclusion of highly divergent outgroup taxa allows us to examine the effects of phylogenetic divergence on exon capture performance. Specifically, we examine the relationship between phylogenetic distance on exon capture sensitivity, specificity, and the proportion of missing data in the final sequence alignments. We also examine the effects of library pool size during multiplexed captures on raw data yield, sequencing depth, and read duplication levels.

## Methods

### Transcriptome Sequencing and Analysis

Four species of hyperoliids representing multiple divergent clades were chosen for transcriptome sequencing: *Afrixalus paradorsalis* (CAS 255487), *Hyperolius balfouri* (CAS 253644), *Hyperolius riggenbachi* (CAS 253658), and *Kassina decorata* (CAS 253990). Whole RNA from a portion of liver sample preserved in RNA Later was extracted using the RNeasy Protect Mini Kit (Qiagen). Samples were evaluated using a BioAnalyzer 2100 RNA Pico chip (Agilent), with RIN scores of 7.0, 7.0, 7.4, and 5.5, respectively. Sequencing libraries were prepared using half reactions of the TruSeq RNA Library Preparation Kit V2 (Illumina), beginning with Poly-A selection for samples with high RIN scores (> 7.0) and Ribo-Zero Magnetic Gold (Epicentre) ribosomal RNA removal for samples with low RIN scores (< 7.0). Libraries were pooled and sequenced on an Illumina HiSeq2500 with 100 bp paired-end reads. Transcriptomic data were cleaned following Singhal (2013). Cleaned data were assembled using trinity (Grabherr et al. 2011) and annotated with *Xenopus tropicalis* (Ensembl) as a reference genome using reciprocal blastx (Altschul et al. 1997) and exonerate (Slater & Birney 2005). We then compared annotated transcripts from the four species to search for orthologs via blast (Altschul et al. 1990). We removed mitochondrial loci from the transcripts. We only kept transcripts with a GC between 40%-70% because extreme GC content causes a reduced capture efficiency for the targets (Bi et al. 2012). Orthologous transcripts with a minimum length of 500 base pairs (bp) were identified across all four samples, resulting in the identification of 2,444 shared transcripts. Transcripts exceeding 850 bp were arbitrarily trimmed to this length for probe design, reflecting a trade-off decision between locus length and the total number of loci included in the experiment. The average pairwise divergence across transcripts among all four samples ranged from 1.4% to 25.9%.

*Availability of Transcriptome Tools.* All the bioinformatics pipelines for transcriptome data processing and annotation are available at https://github.com/CGRL-QB3-UCBerkeley/DenovoTranscriptome.Pipelines for marker development are available at https://github.com/CGRL-QB3-UCBerkeley/MarkerDevelopmentPylogenomics.

### Sequence-Capture Probe Design

The orthologous transcripts were subjected to additional filtering steps before a final gene set was chosen. The initial filtering step applied upper and lower limits on average transcript divergence, eliminating loci with low variation (< 5.0% average divergence) and exceptionally high variation (> 15.0% average divergence), resulting in the removal of 266 genes. The remaining 2,178 genes were examined for repetitive elements, short repeats, and low complexity regions, which are problematic for probe design and capture. The four sets of transcripts per gene (totaling 8,712 sequences) were screened using the repeatmasker Web Server (Smit et al. 2015). This step resulted in the masking of repetitive elements or low complexity regions in 929 sequences, with 7,783 sequences passing the filters. To be conservative, if any of the four transcripts for a gene contained masked sites, that gene was removed from the final marker set, which resulted in the removal of an additional 468 markers. From this reduced set of 1,710 markers, 400 markers with the highest divergence were selected (average divergence ranging from 10.4% to 14.9%) followed by 860 randomly drawn markers from the remaining subset. This marker set was supplemented with five positive controls, which consisted of nuclear sequence data generated using Sanger sequencing for five loci: *POMC* (624 bp), *RAG-1* (777 bp), *TYR* (573 bp), *FICD* (524 bp), and *KIAA2013* (540 bp). The final marker set selected for probe design included 1,265 genes from four species and 5,060 individual sequences.

The final filtered gene set was used to design a MYaits-3 custom bait library (MYcroarray), which consists of 60,060 unique probes per reaction and a total of 48 capture reactions. Following the manufacturers recommendation for capturing sequences of species greater than 5% divergent, 120mer baits were selected, rather than 100mer or 80mer baits. For each locus, we included a sequence from each of the four species; the 5,060 sequences included for probe design totaled 3,983,022 bp, which is approximately 995,700 bp for each full set of loci per species. Following a 2x tiling scheme (every 60 bp) resulted in 60,179 unique baits, therefore 119 probes were randomly dropped to achieve the probe limit.

### Genomic Library Preparation and Pooling

Genomic DNA was extracted from 264 samples (254 ingroup samples, 10 outgroups) using a high-salt extraction method (modified from Aljanabi and Martinez 1997). The DNA was quantified by Qubit DNA BR assay (Life Technologies) and 1700 ng total DNA was diluted in 110 μl of ultrapure H_2_O. A Bioruptor UCD-200 (Diagenode) was used to sonicate the samples on a low setting for 15 minutes, using 30s on/30s off cycling. For each sonicated sample, 4.5 μl of product was run on a 1% gel at 135V for 35 min to ensure fragments were appropriately sized (100–500 bp, average 200–300 bp). Individual genomic libraries were prepared following Meyer and Kircher (2010), with slight modifications, including the use of at least 1600 ng total DNA for library preparation (rather than 500 ng) to remedy the possibility of decreased library diversity resulting from the larger genome size of frogs. We used 7 cycles of post-adapter ligation PCR to enrich the libraries and incorporate a 7bp P7 index, allowing the combination of up to 96 samples in the same sequencing lane. The resulting 50 μl of amplified library product had an average concentration of 35 ng/μl measured by a Nanodrop 1000 spectrophotometer (Thermo Scientific), producing an average yield of 1,750 ng total library DNA.

Samples were pooled for capture reactions according to phylogenetic relatedness as determined by 16S mtDNA data (Portik, unpublished data). Typical pools contained 5–6 genomic libraries, but ranged from 1–7 libraries. All pools contained 1500 ng of total starting DNA, divided equally among the samples included in the pool.

### Sequence Capture Reactions

MYbaits capture reactions were performed following the v2.3.1 manual with some modifications. For each capture reaction library master mix, the pooled libraries were vacuum dried at 45°C for 70 min and re-suspended in ultrapure H_2_O, then combined with 1.66 μl each of human, mouse, and chicken COT-1, and choice of blocking oligos. The combined volume of water for DNA resuspension and volume of blocking oligonucleotides totaled 6.5 μl. An initial three capture reactions were performed on the same library pool to assess the performance of three different types of oligonucleotide blockers designed to anneal to the library adapters during hybridization and prevent daisy-chaining. These blockers consist of the universal blocking oligos (included with the MYbaits kit) which use inosine to block the 7bp index sequence, short blocking oligos which leave the index sequence unblocked, and xGEN blocking oligos (Integrated DNA Technologies), which use proprietary modifications to block the index. Their performance was compared using qPCR analysis of amplified post-capture products, examining enrichment of positive controls and depletion of negative controls. The xGEN blocking oligos performed significantly better in these tests (see Results); we assumed this assessment was a good proxy for sequencing results and these blocking oligos were used for all subsequent capture reactions.

Beyond the slight modifications to the hybridization reaction components discussed above, we followed the manufacturer’s protocol as written, and the hybridization reaction proceeded at 65°C for 27 hours. Individual capture reactions were purified using streptavidin-coated magnetic beads and post-capture products were PCR amplified using four independent reactions of 14 cycles each. These reactions were resuspended in 12 μl of ultrapure H_2_O, and had an average concentration of 15 ng/μl, as measured by Nanodrop. Purified PCR products from the same capture were combined and quantified using a BioAnalyzer 2100 DNA 1000 chip. The combined post-capture amplified products were on average 3.7 ng/μl (range of averages: 1.1–7.2 ng/μl) and the average product size was 398 bp (range of averages: 361–466 bp). Results from Qubit assay were similar, with an average concentration of 4.5 ng/μl (range: 1.0–7.7 ng/μl) for combined post-capture amplified products. The combined post-capture libraries were grouped into three sets (totaling 74, 91, and 92 libraries), pooled in equimolar amounts, and sequenced on three lanes of an Illumina HiSeq2500 with 100 bp paired-end reads.

### Sequence Capture Data Processing

Raw sequence data were cleaned following Singhal (2013) and Bi et al. (2012). In brief, raw fastq reads were filtered using trimmomatic (Bolger et al. 2014) and cutadapt (Martin 2011) to trim adapter contaminations and low quality reads. bowtie2 (Langmead & Salzberg 2012) was used to align the data to *Escherichia coli* (NCBI: 48994873) to remove potential bacteria contamination. We eliminated exact duplicates as well as low complexity sequences using a custom script. Overlapping paired reads were also merged using flash (Magoč & Salzberg) and cope (Liu et al. 2012) to avoid inflated coverage estimate in the overlapping region. The resulting cleaned reads of each individual specimen were *de novo* assembled using abyss (Simpson et al. 2009). We first generated individual raw assemblies using a wide range k-mers (21, 31, 41, 51, 61 and 71) and then used cd-hit-est (Li & Godzik 2006), blat (Kent 2002), and cap3 (Huang & Madan 1999) to cluster and merge all raw assemblies into final, less-redundant assemblies. We used blastn (evalue cutoff = 1e-10, similarity cutoff = 70) to compare the 5,060 target sequences with the raw assemblies of each individual in order to identify the set of contigs that were associated with targets (in-target assemblies). We also ran a self-blastn (evalue cutoff=1e-20) to compare the assemblies against themselves to mask any regions from a contig that matched other regions from other contigs. For each matched contig we used exonerate (Slater & Birney 2005) to define protein-coding and flanking regions. We retained flanking sequences if they were within 500 bp of a coding region. Finally, all discrete contigs that were derived from the same reference transcript were joined together with Ns based on their relative blast hit positions to the reference. Most of the final in-target assemblies contain multiple contigs, and each includes both coding regions and flanking sequences if captured.

Cleaned sequence data were then aligned to the resulting individual-specific intarget assemblies using novoalign (Li & Durbin 2009) and we only retained reads that mapped uniquely to the reference. We used Picard (http://broadinstitute.github.io/picard/) and gatk (McKenna et al. 2010) to perform realignment. We finally used samtools/bcftools (Li et al. 2009) to generate individual consensus sequences by calling genotypes and incorporate ambiguous sites in the intarget assemblies. We kept a consensus base only when the site depth is above 5X. We masked sites within 5 bp window around an indel. We also filtered out sites where more than two alleles were called. We converted FASTQ to FASTA using seqtk (https://github.com/lh3/seqtk) and masked putative repetitive elements and short repeats using repeatmasker (Smit et al. 2015) with vertebrata metazoa as a database. We removed markers if more than 80% of the bases were Ns. We then calculated read depth of each individual marker and filtered out loci if the depth fell outside 1^st^ and 99^th^ percentile of the statistics. We also eliminated markers if the individual heteozygosity fell outside the 99^th^ percentile of the statistics. The final filtered assemblies of each individual specimen were aligned using mafft (KAtoh & Standley 2013). Alignments were then trimmed using trimal (Capella-Gutierrez et al. 2009). We removed alignments if more than 30% missing data (Ns) are present in 30% of the samples. We also removed alignments if the proportion of shared polymorphic sites in any locus is greater than 20%.

*Availability of Sequence Capture Data Tools.* The bioinformatic pipelines of sequence capture data processing are available at https://github.com/CGRL-QB3-UCBerkeley/denovoTargetCapturePhylogenomics.

### Sequence Capture Efficiency Evaluation

*Sequencing Depth, Duplication Levels, and Pooling Sizes.* To evaluate capture efficiency, average per-base sequence depth, or coverage, was calculated separately for the exon sequences and for the flanking sequences of each sample. The coverage at each base pair site for either data set was inferred using the samtools (Li et al. 2009). The per base pair coverage estimates for all sequences (exon or flanking) associated with each transcript (up to 1,260 genes) were averaged, resulting in a set of average coverage estimates across loci. The resulting output of the set of average coverage estimates was used to infer the median, upper and lower quartiles, and range of coverage estimates using samples or genes as factors. These calculations were performed and automated across samples using python scripts and the output was visualized in R. Differences in the levels of coverage were examined using pooling size as a factor. To control for differences in coverage possibly resulting from phylogenetic distance, comparisons were only made among pools of the ingroup genus *Hyperolius* (160 samples, 28 captures).

Duplication refers to the number of non-unique sequencing reads, which were eliminated from our sequence capture data processing pipeline. The level of duplication among reads, expressed as a percentage, was estimated by dividing the number of duplicate reads by the total number of raw reads. Differences in the levels of duplication were examined using pooling size as a factor, compared across the genus *Hyperolius*. The amount of raw data (in bases) was also compared across pool sizes using the genus *Hyperolius.*

*Sequence Capture Sensitivity.* Sensitivity refers to the percentage of bases of target sequences that are covered by at least one read, and here the target refers to the exons of each gene. To calculate this metric, the final in-target assemblies (including exons and flanking sequences) of each sample were compared to a set of transcript sequences used for probe design, from only one of the design species, using BLASTN with a evalue cutoff of 1e-10. This was automated using custom scripts to produce output files of all blast hits for each sample. For each output file, any overlapping base pair coordinates for blast hits within a locus were merged. Following the merging of coordinates, the number of base pairs for all exon blast hits per locus was totaled, and was divided by the total length of the transcript sequence to calculate the sensitivity per transcript. The total number of base pairs from all exon blast hits was divided by the total number of base pairs of all the transcript sequences, producing an overall estimate of sensitivity per sample.

*Sequence Capture Specificity*. Specificity is a metric that measures how many base pairs of cleaned reads are aligned to target sequences, expressed as a percentage. In this experiment, the target sequences are represented in two ways: in-target assemblies (exons and their associated flanking sequences), and exons only. For each sample, bam files were converted to sam file format using the samtools view function and the total number of base pairs aligned within the exon sequences and flanking sequences were counted by parsing the bam files. To estimate base pairs aligned with transcript exon sequences only, the sample bam file was converted to sam format using the associated bed file containing base pair coordinates for exons only, and total aligned base pairs were calculated in the same manner. The number of cleaned read base pairs was calculated from the summing the read lengths contained within cleaned reads files.

*Exon Coverage Uniformity.* The uniformity of coverage across the length of exons was examined using both longer (> 200 bp) and shorter (61–100 bp) exons. Exons matching these criteria were filtered out from bed files containing exon coordinates independently for each sample. For longer exons, five bins of 10 bp increments were created for both the 5’ and 3’ ends, resulting in the generation of ten additional bed files per sample. For shorter exons, three bins of 10 bp increments were created for both the 5’ and 3’ ends, resulting in the generation of six additional bed files per sample. Each bed file was used to calculate the per base pair coverage for a specific end bin using samtools. These per base pair coverage values were averaged within exons, and all averages of exons for a particular bin were subsequently combined across 50 randomly chosen samples. The values across bins were visualized in R to assess the median, upper and lower quartiles, and range of coverage estimates.

*Effects of Phylogenetic Distance.* We sought to test the relationship between phylogenetic distances and several evaluation metrics to determine if Sanger sequence data have predictive power for exon capture success. Phylogenetic distance was calculated as the average of uncorrected pairwise differences between samples and the four design species. These divergence estimates were calculated using the five positive controls (nuclear loci from Sanger sequencing). As this information would generally be available to researchers before designing such an experiment, these loci provide an a priori estimation of divergence across the focal group. The effects of phylogenetic distances on capture specificity, sensitivity, and duplication were investigated using simple linear regressions. The values for the above metrics were averaged for each genus, providing values for a total of 23 genera for comparison. Average phylogenetic distances ranged from 6.7–18.3%, representing divergences up to 103 million years old (Portik & Blackburn, in prep).

*Evaluation of Exon Phylogenetic Informativeness.* The resulting alignments of exon-only data and flanking region data were evaluated for taxon number, sequence length, percentage of missing data, and proportion of informative sites. These results were visualized in R, and the relationship between the number of informative sites and alignment length was investigated using a simple linear regression. The relationship between phylogenetic distance and missing data was also investigated using a simple linear regression. The percentage of missing data was calculated from the final concatenated alignment of exon-only loci that passed multiple post-processing filters, including a minimum length of 90 bp, no more than 80% missing data per sequence in alignments, and no more than 30% total missing data across an alignment. These filters were enforced using a custom alignment refinement python script for all alignments.

*Availability of bioinformatics tools.* All custom python scripts for sequence capture performance evaluation are available on github (https://github.com/dportik/). These include tools for automating the calculation of coverage, duplication, sensitivity, specificity, and coverage uniformity. Additional scripts are available for evaluating and refining DNA sequence alignments.

## Results

### Effects of Blocking oligos

Quantitiative PCR reactions were performed for a positive control nuclear locus (KIAA2013) targeted by the hybridization probes and a negative control nuclear locus (49065) not targeted by the capture probes. All reactions were standardized for the same input amount of DNA (4ng). For the positive control, all post-capture curves show an expected leftward shift relative to the pre-capture, indicating that the concentration of copies of the KIAA2013 locus has increased significantly in the post-capture library pools. Of the three blocker types, the greatest change in enrichment is observed with the post-capture pool using xGen blocking oligos (11.9 cycle shift), rather than the universal blocking oligos or short blocking oligos (10.3 cycle shifts) (Fig. 1). For the negative control, all post-capture curves show an expected rightward shift relative to the pre-capture, indicating that the concentration of copies of the 49065 locus has decreased in the post-capture library pools. However, the universal blocking oligos and short blocking oligos show only minor differences from the pre-capture library (1.9 and 1.0 cycle shifts respectively) (Fig. 1). In contrast, the post-capture pool using xGen blocking oligos has shifted considerably (10.3 cycle shift), indicating that non-target regions have been significantly depleted from the library pool (Fig. 1). This is reflected in the post-capture library quantification, in which higher amounts of DNA were detected in the universal blocker reaction (23 ng/μL) and short blocker reaction (23.7 ng/μL), compared to the xGen reaction (14 ng/μL), indicating that more non-targeted sequences were accidentally captured during the hybridization.

**Figure 1.**
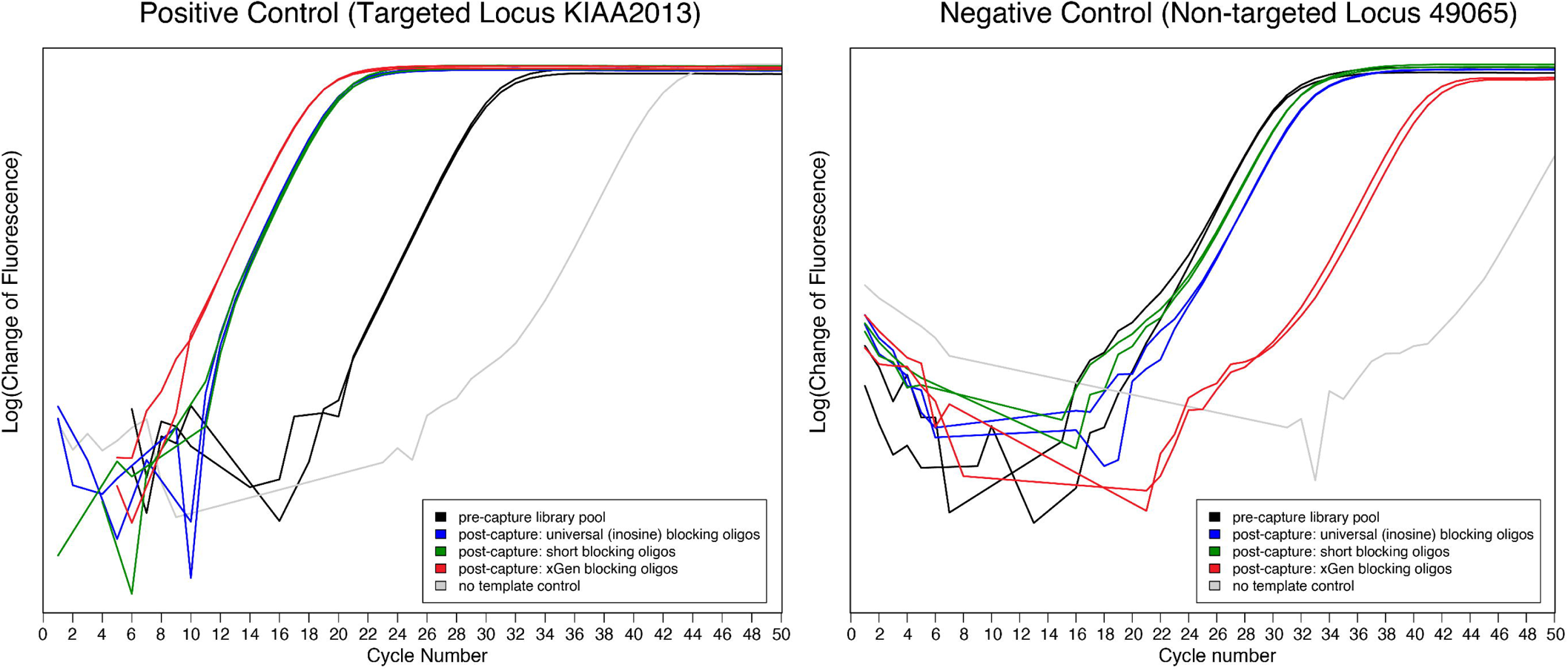
Quantitiative PCR plots for the positive control nuclear locus (KIAA2013) and the negative control nuclear locus (49065). Assessment of the relative success of the capture can be made by comparing the position of the curves of the library pool prior to capture (black) to the three curves produced by library pools after capture using different blocking oligo strategies (blue, green, red). The largest cycle shifts in both enrichment and depletion occur using the xGen blocking oligos (red), with substantially better performance occurring for the depletion of non-targeted sequences.

### Sequence Capture Data

The average number of reads sequenced across the 264 samples is 2,422,484 (range: 415,439–6,899,259 reads), and as we sequenced 100 bp paired-end reads, the average total base pair yield is 484.4 Mb (range: 83.0–1,379.8 Mb). In addition to the removal of low complexity and low quality reads, the raw reads were filtered to remove exact duplicates, adapters, and bacteria contamination. After these filtration steps, the average number of base pairs of cleaned reads was 331.9 Mb (range: 65.3–789.6 Mb); on average 68% of the raw base pairs passed the quality control filters.

The assemblies were assigned to targeted transcripts, and the resulting in-target assemblies contained a mix of exon sequences and non-coding flanking sequences (Fig. 2A). The length of the in-target assemblies were often several thousand base pairs, much longer than the original targeted transcript sequences (which were maximally 850 bp), illustrating a significant amount of non-coding flanking sequence data associated with each exon was captured (Fig. 2A). By trimming the flanking sequences, the concatenated exons closely match the original transcript sequence lengths (Fig. 2B). Across all targeted loci and samples, the median number of exons per transcript is four, but ranged from a single exon to eleven exons per transcript (Fig. 3). The average length of exons within transcripts recovered is 153 bp, but the data set revealed a wide range in sizes, from shorter exons (< 100 bp) to longer exons (> 600 bp) that cover the entire transcript sequence used (Fig. 4).

**Figure 2.**
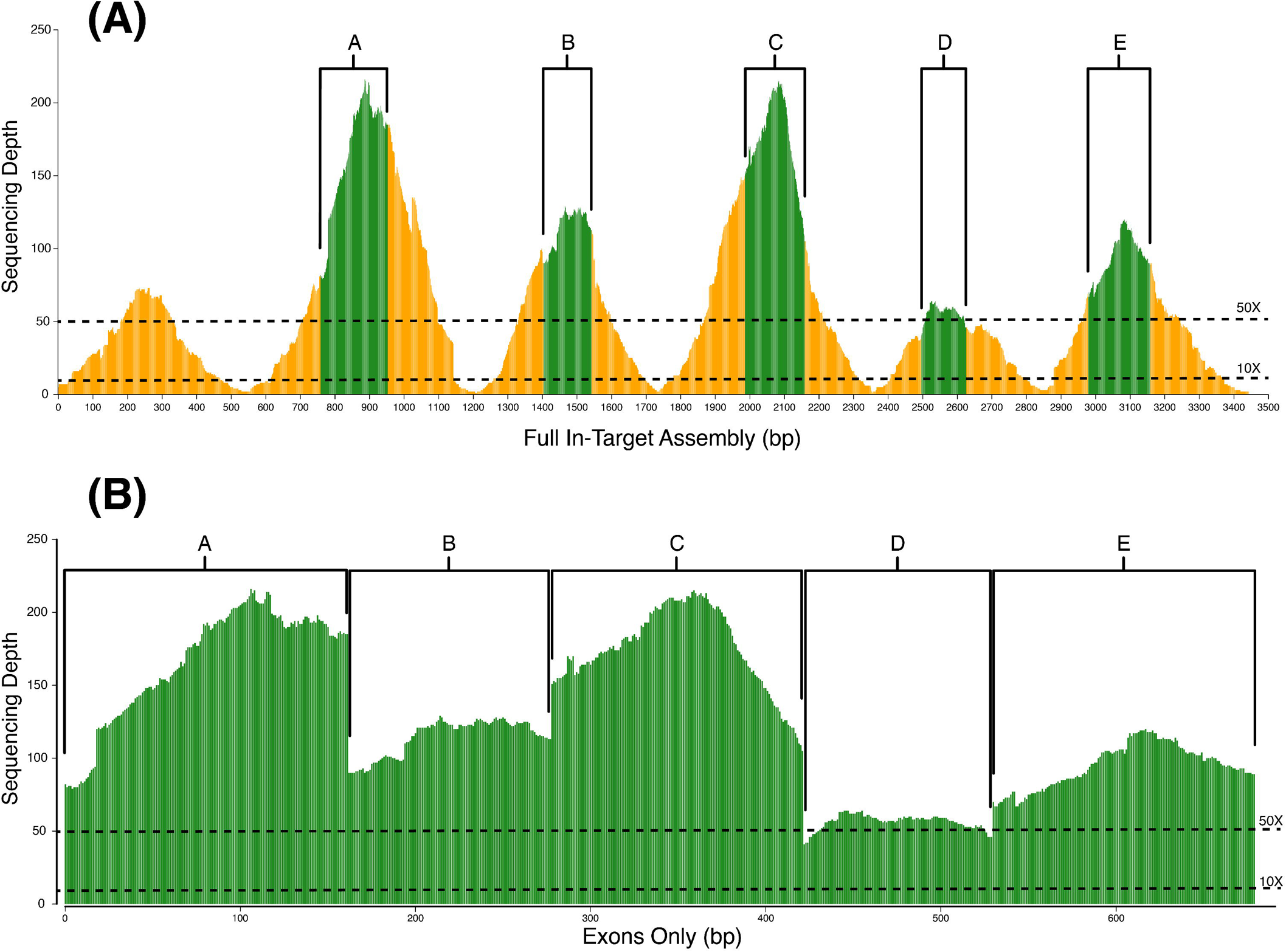
An example sequencing depth (coverage) plot for (A) an in-target assembly and (B) exon-only contig of the same locus from one sample. Exons matching the transcript sequence used for probe design are colored green and labeled (A–E, matching in both plots), and non-coding flanking regions are colored orange. Both 50X and 10X coverage levels are indicated by dotted lines.

**Figure 3.**
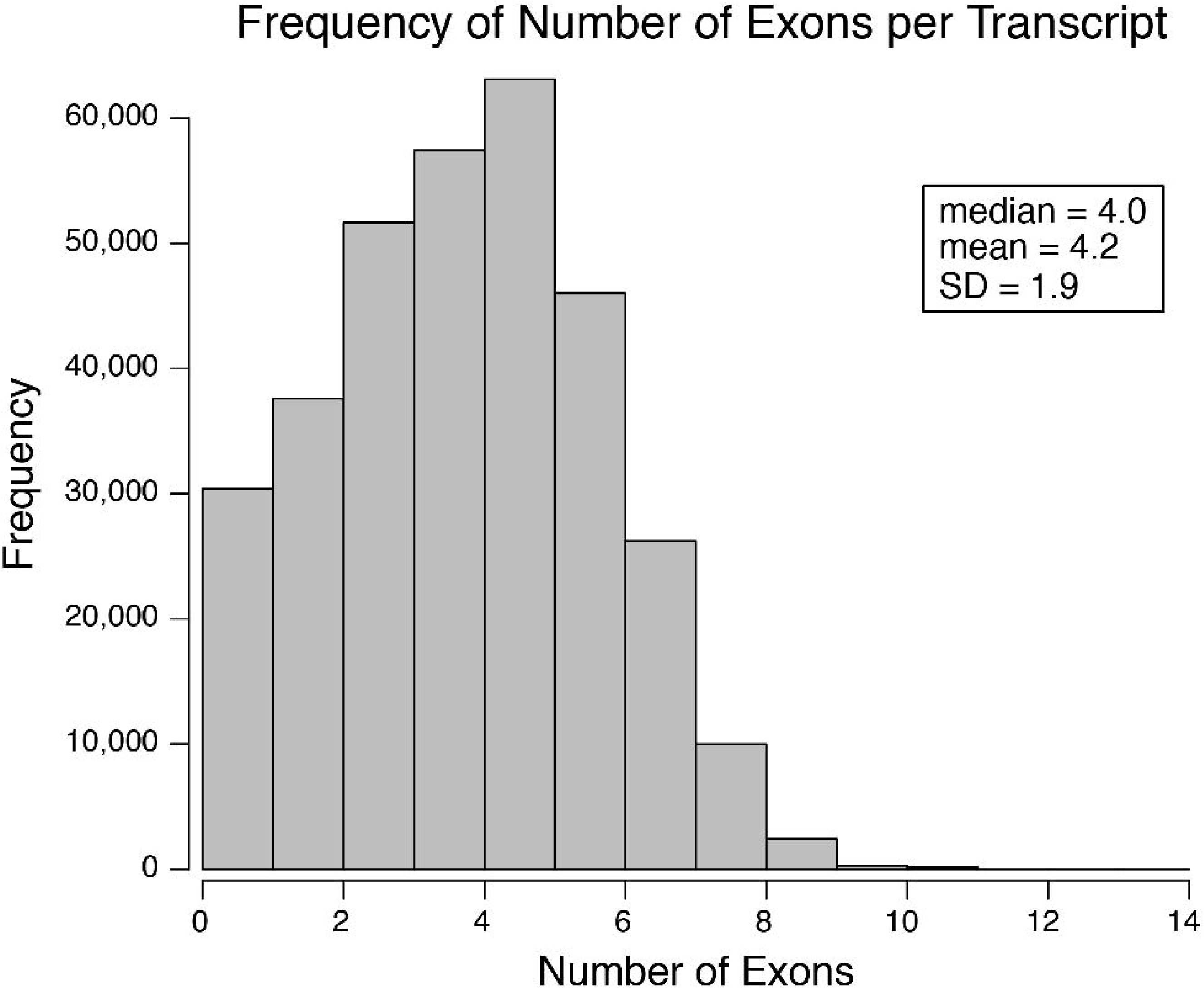
A frequency distribution for the number of exons detected in the fully assembled and merged contigs across all samples. The median number of exons per transcript is four.

**Figure 4.**
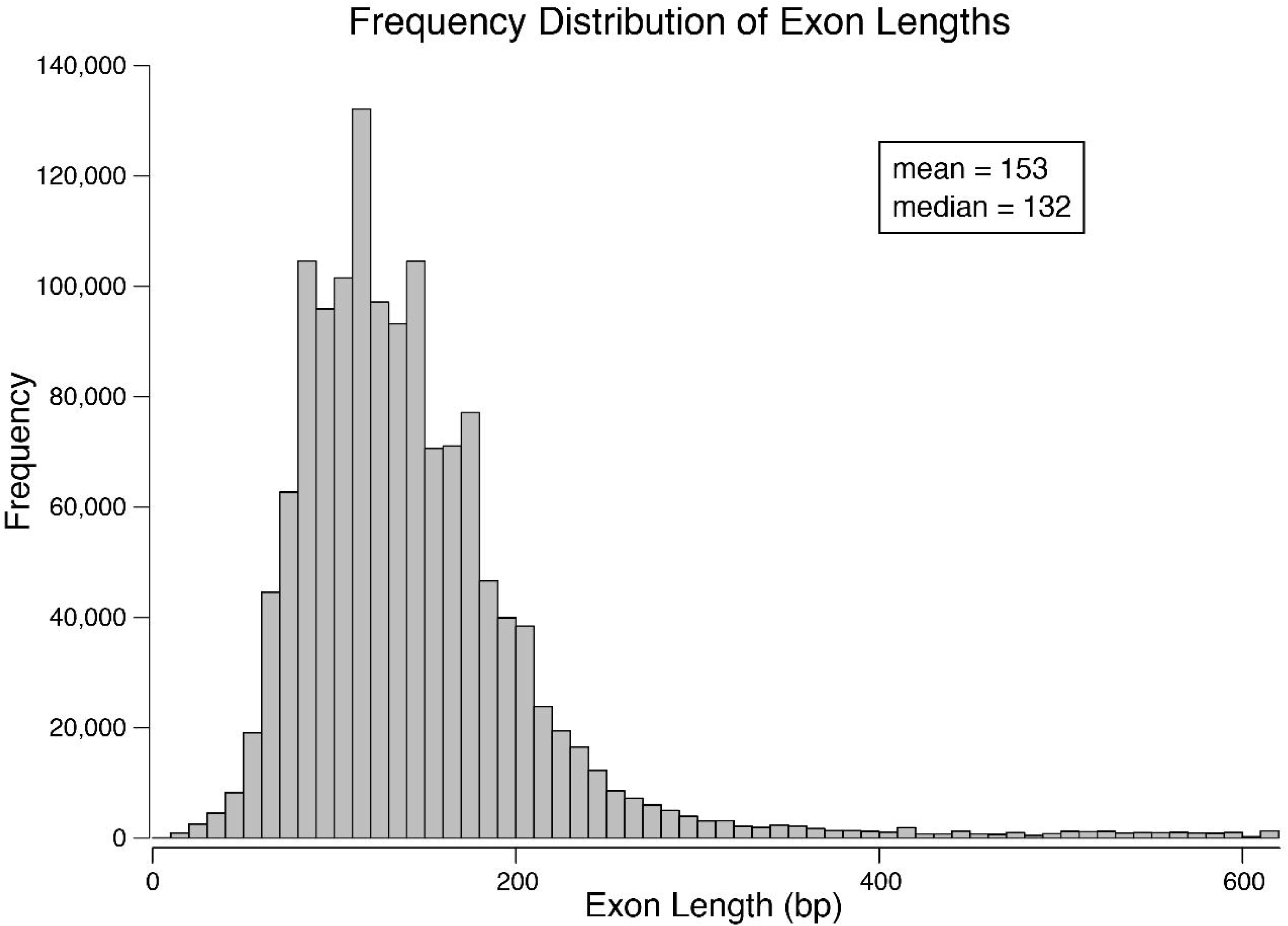
A frequency distribution for the length of each exon detected within a fully assembled and merged contig, across all samples. The average length is 153 bp, and the median length is 132 bp.

### Sequencing Depth and Duplication Levels

The sequencing depths of merged contigs showed variation between loci and across samples, but the most pronounced differences in coverage occurred between the exon and flanking regions (Fig. 2A). The average sequencing depth across all exons for all samples averaged 142.4X (n = 1,372,603 exons), whereas the flanking regions averaged 45.5X. This result is consistent with expectations for transcriptome-based exon capture, as the probe design only considered exon regions. Despite not specifically targeting these adjacent non-coding regions, this experiment clearly demonstrates non-coding regions can be captured and sequenced with sufficiently high coverage. Because the estimates of sequencing depth only consider sites that are captured, relating coverage to phylogenetic distance is not a meaningful metric. We did consider the effect of pooling size on coverage, but within a single genus that was the main focus of the experiment and for which capture results were very similar (genus *Hyperolius*). A comparison of pool sizes (1–2, 4–7) revealed no significant differences in sequencing depth across all loci based on the student’s t-test (Fig. 5). Similarly, there do not appear to be to be differences in raw data yield (in total base pairs) for different pool sizes (Fig. 5), though low sample sizes in smaller pools preclude rigorous testing.

**Figure 5.**
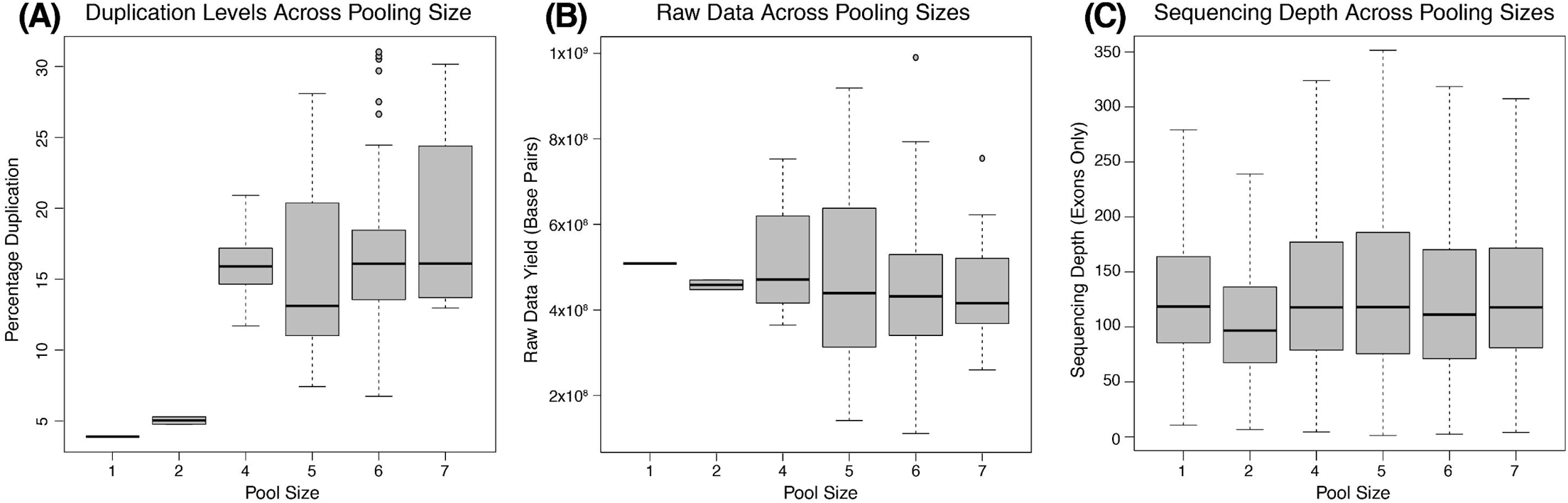
Boxplots of pooling sizes and (A) duplication levels, (B) raw data yield, and (C) sequencing depth. The boxplots depict the median, upper and lower quartiles, and range for each metric.

The duplication levels among reads are an indicator of the diversity of sequences captured, with high duplication implying a less diverse post-capture library relative to post-capture libraries with lower duplication levels. In general, a higher number of post-capture PCR cycles are expected to produce higher levels of duplication among samples. In this experiment, all post-capture PCR reactions used the same number of cycles; therefore, our comparison of duplication levels is an indicator of post-capture sequence diversity rather than a methodological artifact. Levels of duplication were similar between the ingroup (average: 17.2%, range: 3.4%–37.9%) and outgroups (average: 16.5%, range: 5.0%–24.8%). We tested for a relationship between duplication level and phylogenetic distance using a simple linear regression, and found the regression was not significant (F(1, 21) = 0.79, p = 0.38). Though phylogenetic relatedness may not be a predictor of duplication levels, there is a clear pattern of differences in duplication levels across pooling sizes (Fig. 5). Pools with a single individual or two individuals have much lower duplication levels (3.9% and 5.1%, respectively) than pools with four or more individuals (range of averages: 15.7%–19.0%) (Fig. 5). Small samples sizes precluded statistical testing for these differences between smaller and larger pools, but these data indicate pooling size is much more likely to affect duplication levels than other factors such as phylogenetic distance. We did not find significant differences in duplication levels between larger pools, and this strongly suggests pooling seven individuals did not negatively impact the resulting diversity of sequences captured among samples. Additional replicate captures of larger pool sizes can help determine at which point captured sequence diversity is impacted and establish limitations in pooling sizes for successful capture.

### Exon Coverage Uniformity

Using 50 random ingroup samples, sequencing depth values were calculated for the edges of exons in 10 base pair bins, with 5 bins included for longer exons (> 200 bp) and 3 bins included for shorter exons (61–100 bp). At the 5’ and 3’ ends of longer exons, the average coverage is 117.2X and 124.0X, respectively (Fig. 6, Table 1). These values increase slightly across bins towards the center, with both the 5’ and 3’ 41–50 bp bins having approximately 165X coverage (Fig. 6, Table 1). The coverage values for edge bins of shorter exons were lower, but in general showed the same trend increasing towards the center (Fig. 7, Table 2). Here, the average coverage of the 5’ and 3’ ends is 74.9X and 80.4X, respectively, with the most central bins (21–30 bp) exhibiting 83.7X and 86.2X coverage (Fig. 7, Table 2). Together, these results indicate high uniformity in sequencing depth across the length of short exons, and demonstrate only a slight decrease in coverage towards the edges of longer exons.

**Figure 6.**
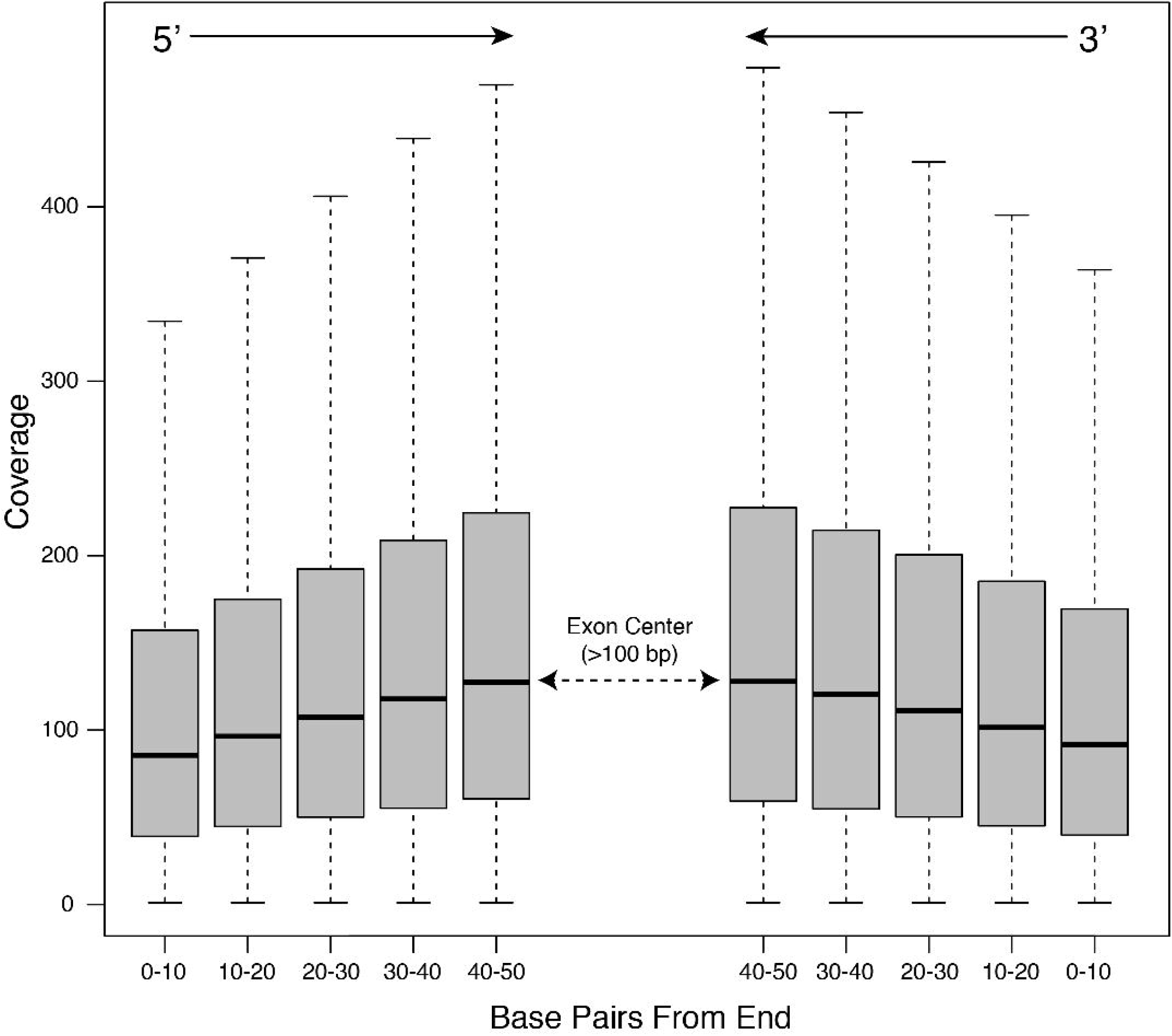
The average binned coverage of the edges of long exons (> 200 base pairs). Bins are in 10 base pair increments, with five bins on the 5’ and 3’ ends. Estimates are based on 50 randomly chosen ingroup samples.

**Figure 7.**
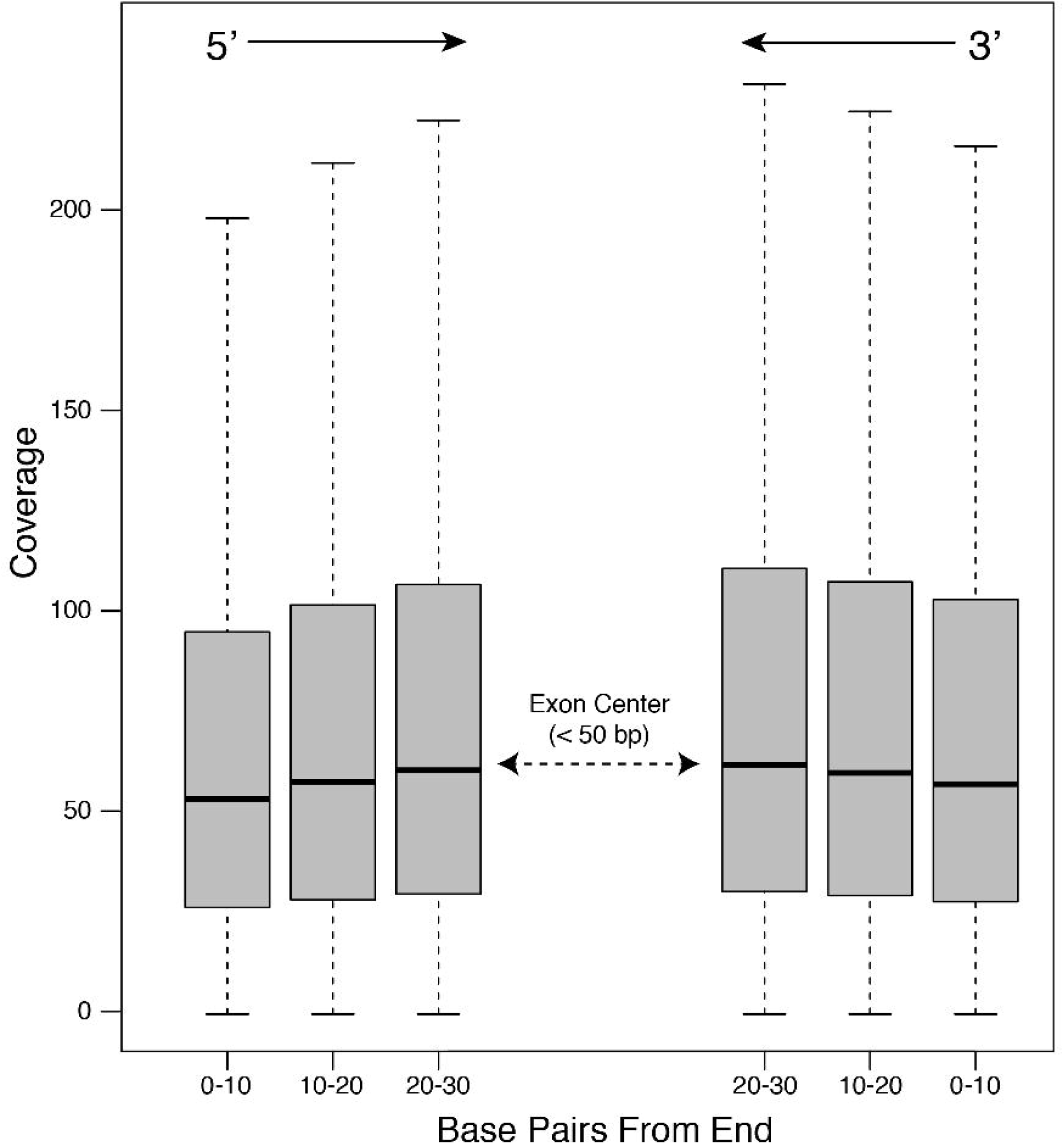
The average binned coverage of the edges of short exons (61–100 base pairs). Bins are in 10 base pair increments, with three bins on the 5’ and 3’ ends. Estimates are based on 50 randomly chosen ingroup samples.

### Sensitivity, Specificity, and the Effects of Phylogenetic Divergence

We explored capture sensitivity, the percentage of bases of in-target assemblies that are covered by at least one read, across all samples in our experiment. In general, sensitivity varied between genera but was relatively consistent within genera (Fig. 8A). The average across all ingroup samples is 80.1% (range 52.1%–91.8%), whereas outgroup samples averaged 33.8% (range 20.7%–42.2%). A simple linear regression was calculated to predict sensitivity (%) based on phylogenetic distance. A significant regression equation was found (F(1, 21) = 79.58, p < 0.001), with an adjusted R^2^ of 0.78 (Fig. 9). Sensitivity is equal to [108.19 + −4.57*(average pairwise divergence)] percent when pairwise divergence is measured as a percent; sensitivity decreased 4.57% for each percent increase of pairwise divergence.

**Figure 8.**
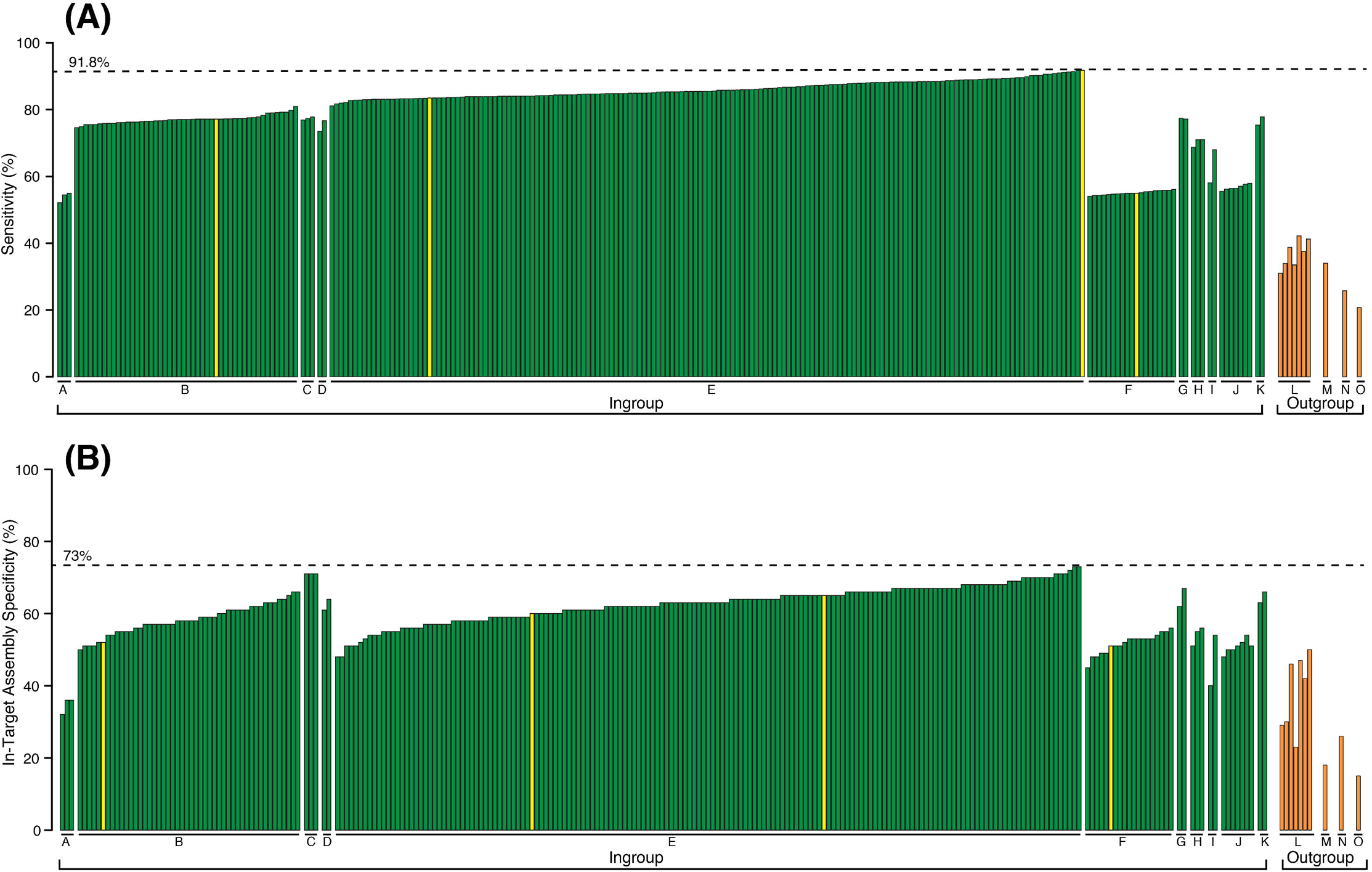
Barplot of (A) sensitivity and (B) specificity, across all samples. Labels A-K refer to ingroup genera denoted by green (A: *Acanthixalus,* B: *Afrixalus,* C: *Cryptothylax,* D: *Heterixalus,* E: *Hyperolius,* F: *Kassina,* G: *Morerella,* H: *Opisthothylax,* I: *Paracassina,* J: *Phlyctimantis,* K: *Tachycnemis*) and labels L–O refer to outgroups denoted by orange (L: Arthroleptidae, M: Brevicipitidae, N: Hemisotidae, O: Microhylidae). Yellow indicates the species used for transcriptome sequencing and probe design.

Specificity is a metric similar to sensitivity, but it measures the percentage of base pairs of cleaned reads that can be aligned to target sequences. We investigated specificity using the in-target assemblies (exons and flanking regions) and exons only. Specificity varied among genera (Fig. 8B), and across all ingroup samples averaged 60.2% (range 32.0%–73.0%), whereas outgroup samples averaged 35.6% (range 15.0%–50.0%). As expected, specificity of the exon only data set was lower, and ingroup genera exhibited higher specificity (47.3%, range: 26.0%–65.0%) than outgroup genera (27.7%, range: 13.0%–40.0%). Using specificity results from the in-target assemblies, a simple linear regression was calculated to predict specificity (%) based on phylogenetic distance. A significant regression equation was found (F(1, 21) = 44.1, p < 0.001), with an adjusted R^2^ of 0.66 (Fig. 9). Specificity is equal to 83.99 + −3.26*(average pairwise divergence) percent when pairwise divergence is measured as a percent. Specificity decreased 3.26% for each percent increase of pairwise divergence.

**Figure 9.**
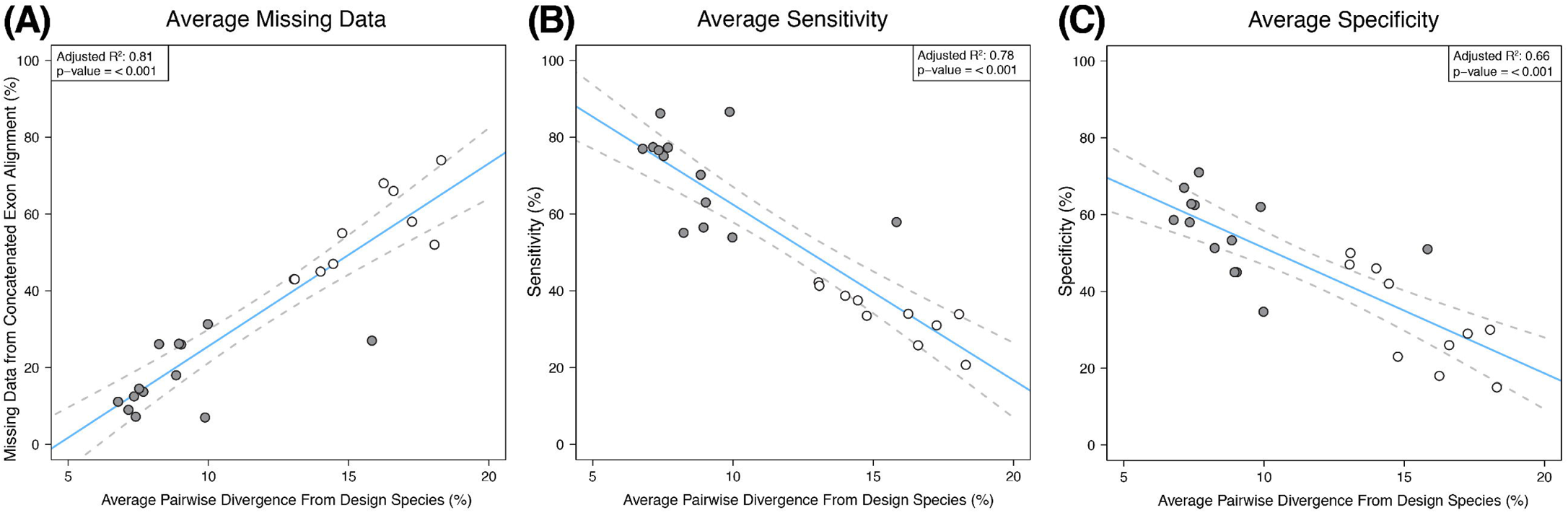
Plots of linear regressions of (A) missing data from the concatenated exon alignment, (B) sensitivity, and (C) specificity, using the average pairwise divergence from probe design species as the independent variable.

### Sequence Alignments and Informativeness

There were a total of 1,047 exon-only alignments and 287 flanking region alignments that passed all filtering criteria. Together, the concatenated alignment of flanking and exon data totals 631,127 base pairs.

For exon-only alignments, the average number of taxa included is 250, average per locus length is 536 bp, average level of missing data is 8.9%, and the average proportion of informative sites is 38.3%. The concatenated alignment of the exon-only loci totals 561,180 base pairs. The average proportion of missing data in the concatenated alignment for the ingroup samples is 10.7% (range 3%–35%), and is 55.1% (range 43%–74%) for the outgroup samples. A simple linear regression was calculated to predict missing data levels in the final exon-only alignments, based on phylogenetic distance. A significant regression equation was found (F(1, 21) = 96.78, p < 0.001), with an adjusted R^2^ of 0.81 (Fig. 9). Missing data is equal to [−22.07 + 4.76*(average pairwise divergence)] percent when pairwise divergence is measured as a percent. Missing data increased 4.76 percent for each percent increase of pairwise divergence. A simple linear regression was also calculated to predict the number of informative sites in an exon-only locus based on the length of the locus. A significant regression equation was found (F(1, 1045) = 5666, p < 0.001), with an adjusted R^2^ of 0.84 (Fig. 10). The number of informative sites is equal to [-1.89 + 0.38*(alignment length)]. Informative sites increased 0.38 base pairs for each base pair increase in alignment length (Fig. 10).

**Figure 10.**
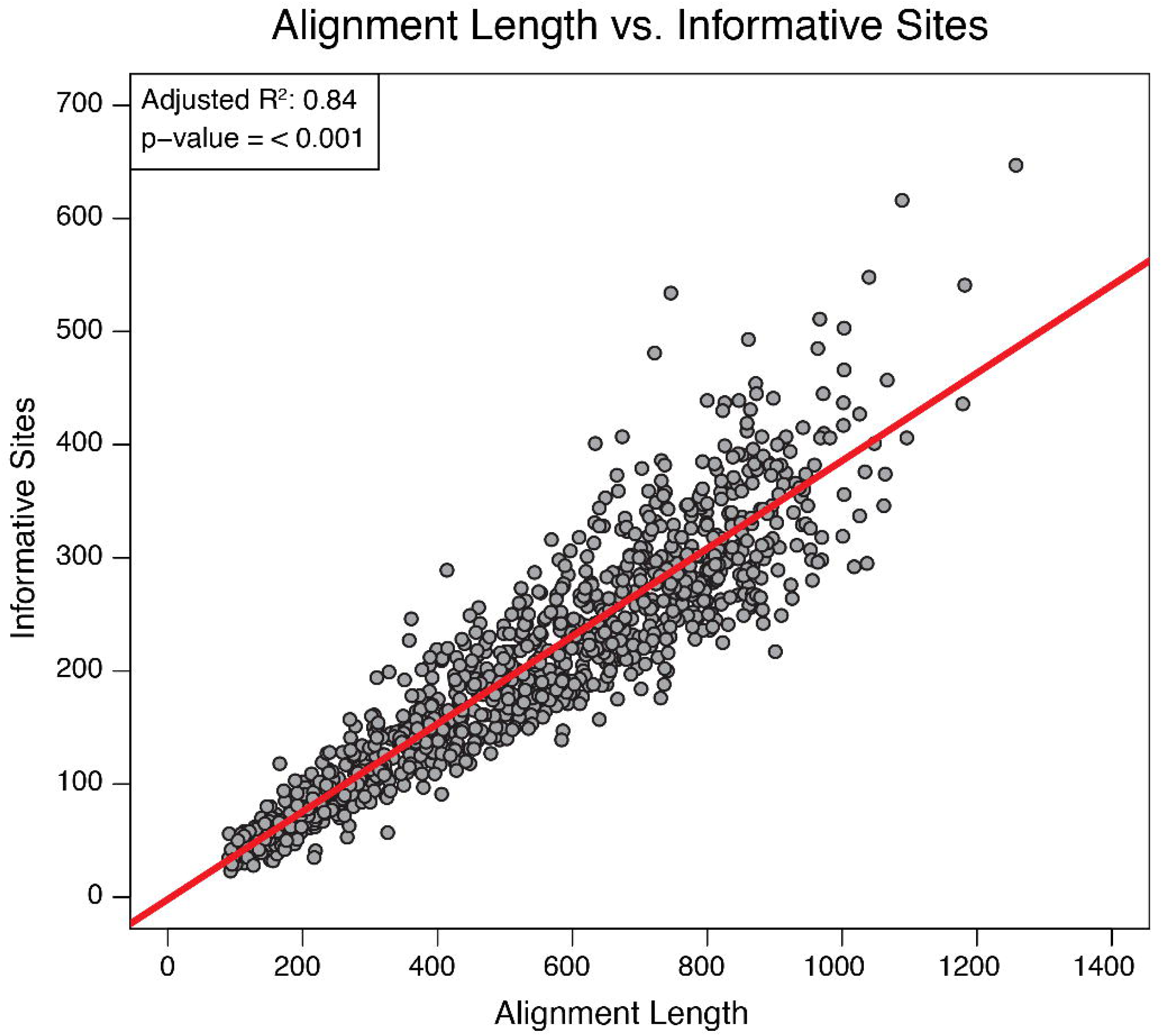
Linear regression of alignment length and the number of informative sites. Each dot represents a unique exon-only alignment, for a total of 1,047 loci. A significant regression equation was found (F(1, 1045) = 5666, p < 0.001), with an adjusted R^2^ of 0.84.

For flanking region alignments, the average number of taxa included is 250, the average length is 243 bp, the average level of missing data is 12.4%, and the average proportion of informative sites is substantially higher than exon-only alignments at 77.4%. The concatenated alignment of the flanking-only loci totals 69,947 base pairs. The average proportion of missing data in the concatenated alignment for the ingroup samples is 15.4% (range 6%–40%), and is 50.6% (range 42%–68%) for the outgroup samples. The non-coding flanking loci are generally more difficult to align, especially as phylogenetic distance increases. For the purpose of this study, we performed alignments across all samples, which is likely to have contributed to lower quality alignments and failure to pass specific missing data filters. We therefore emphasize if flanking sequence alignments are performed for the ingroup only, or even subclades of the ingroup, it should result in more and longer alignments recovered.

## Discussion

We used a custom transcriptome-based exon capture, designed without the use of a reference genome, to successfully generate a large informative phylogenomic data set across divergent lineages of frogs. We accomplished this using transcriptome sequences directly for probe design, resulting in the additional recovery of a significant amount of highly variable non-coding sequence data. We generated sequence alignments for 1,047 of the 1,260 transcriptome-based loci, with an average of 250 (of 264) taxa present per alignment. The combination of exon and flanking region data resulted in a concatenated alignment of 631,127 base pairs. Based on the results of our experiment, we discuss the overall efficiency of capture, results of using transcript sequences for probe design, effects of phylogenetic distance, and recommendations for pooling size and blocking oligos.

The effects of blocking oligos are non-trivial, and have great potential to affect the capture efficiency (Fig. 1). Although the enrichment of target loci is accomplished using short blockers, universal blockers, and xGen blockers, there are critical differences in the level of depletion of non-targeted loci among blockers. The xGen blockers significantly outperformed the short blockers and universal blockers in the depletion of non-targets (Fig. 1). The higher concentration of DNA in post-capture libraries of the universal and short blockers represents a large carry-over of non-targets, which ultimately translates to significant reductions in both sensitivity and specificity and increases in PCR duplication rates. This is particularly important to consider for organisms with larger genome sizes, such as amphibians, which are likely to suffer from reductions in sensitivity and specificity and higher duplication rates based on genome size alone. The cost of xGen blockers is significantly higher per reaction, but may ultimately reduce the amount of sequencing effort required to obtain high quality sequence data. We therefore strongly recommend the testing of blocking oligos using a qPCR assay before conducting the main capture experiment, as the specificity, sensitivity, and duplication levels can be greatly improved with appropriate blocking oligos.

A main question concerning sequence capture is simply how many individuals can be pooled in a reaction, which has important implications for reducing costs and increasing the sampling for a given project. We tested a range of pool sizes (1–7 samples) within the genus *Hyperolius* (160 samples, 28 captures) to determine the effects of pooling on raw data yield, sequencing depth, and duplication levels. We found no differences in raw data yield or sequencing depth across pools, but our results indicate duplication levels vary across pooling sizes (Fig. 5). We demonstrate duplication levels rose from 4–5% in 1–2 sample pools to an average of 15–19% in the 4–7 sample pools. These levels were acceptable for obtaining high quality sequence capture data for our experimental design. We did not detect a significant increase in duplication levels for pools of seven samples, suggesting the upper limit for sample pooling was not reached, though this topic requires additional investigation. Although pooling size does affect PCR duplication levels, we again emphasize that these effects can be strongly exacerbated through the use of less efficient blocking oligos.

Phylogenetic distance has a predictable effect on capture sensitivity, specificity, and the proportion of missing data in the final sequence alignments. As expected, more divergent species experienced drops in sensitivity and specificity, and their proportion of missing data increased (Fig. 9). Though these results are intuitive, our findings our useful in at least two ways. First, we demonstrate that for the most distant outgroup in our experiment (family Microhylidae), which shared a common ancestor with the probe design species 103 million years ago, we recovered 23% of the total exon sequence data (roughly 146,000 bp). Our experiment was focused on sequence capture within a single family, but successful sequence capture occurred for highly divergent outgroup species, albeit with a predictable reduction in efficiency. Second, the regression equations relating capture efficiency metrics to average pairwise divergence can serve as a starting point for other researchers in determining the phylogenetic breadth of their capture experiments. Our comparisons are made using nuclear sequence data generated prior to our experiment. These empirical data, though based on frogs, allow an approximation of the effects of phylogenetic distance on metrics generally used to characterize capture efficiency, and can set realistic expectations for the overall success of sequence capture based on phylogeny. This approximation requires Sanger sequencing only a small number of nuclear loci for a subset of the target group, information that should generally be acquired before beginning a large-scale capture experiment.

Our experimental design used transcriptome sequences of four species from divergent ingroup clades to design capture probes, and we recovered high quality sequence data across the ingroup genera. The use of four sets of sequences for each locus ultimately reduced the total number of loci that were included in our probe design, and the trade-off between number of loci and variability in probe design is important to consider for exon capture design. Unfortunately we cannot assess whether probe sets from certain species were more efficient in capturing sequences, and it is unclear how using a single species would have affected the outcome of this experiment. Using a single species for probe design in our case would have allowed for the inclusion of approximately 5,000 loci, rather than 1,260. A possibility for reducing the number of design species is to reconstruct ancestral sequences for deeper nodes of the ingroup, and use these sequences for probe design. Though there are many options for probe design, our results demonstrate sampling multiple divergent ingroup species is a highly effective strategy for capture across larger phylogenetic scales.

Our experiment tested the direct use of transcriptome sequences for probe design, thereby circumventing the use of reference genomes for identifying intron-exon boundaries to filter out short exons. This approach was highly successful, and we recovered short and long exons with a high uniformity in coverage (Figs. 6, 7) as well as a large quantity of highly variable non-coding flanking regions. The average size of individual exons (153 bp) within loci is close to predictions of average exon lengths across the genome of *Xenopus laevis* (~200 bp), and we found most of the 850 bp transcriptome sequences contained four exons (Figs. 3, 4). We successfully captured large quantities of short exons (< 100 bp) (Fig. 4), a feature that may be appealing for researchers targeting short loci. The probe design spanning intron-exon boundaries did not reduce coverage towards the ends of exons (Figs. 6, 7), and returned thousands of base pairs of non-coding flanking sequences per in-target assembly. The resulting alignments of non-coding regions show high levels of variation, with an average proportion of 77% informative sites (compared to 38% of exon-only regions). These flanking regions can be incorporated into population genetics or phylogenetic analyses, similar to UCE and anchored hybrid enrichment approaches. Our pipeline allows alignments to be made with the full in-target assemblies, exon regions only, or flanking regions only, providing flexibility for decisions about sequence data analysis.

Transcriptome-based exon capture is an effective method for producing large sets of orthologous markers with predictable levels of informativeness in non-model systems. This method can be applied to population level questions by sequencing transcriptomes within a population, or applied to larger phylogenetic scales using the transcriptomes of divergent species. As this approach and other types of sequence capture gain popularity, the reporting of empirical data can enhance the ability of researchers to choose the appropriate capture approach or aid in the design of custom sequence captures. We have outlined our experimental design, including probe design from transcriptome sequences, as well as reaction-specific decisions about blockers and capture pooling schemes. For this type of transcriptome-based exon capture, information about the number of exons in transcripts, their lengths, and the recovery of flanking sequences should be discussed. Finally, efforts to relate any of the above measures to phylogenetic distance would greatly benefit researchers planning a sequence capture experiment for non-model systems.

## Acknowledgments

Lab work conducted by DMP was funded by a National Science Foundation DDIG (DEB: 1311006), the EECG Research Award (American Genetic Association), and by D.C. Blackburn, R.C. Bell, and J.A. McGuire. This work used the Vincent J. Coates Genomics Sequencing Laboratory at UC Berkeley, supported by NIH S10 Instrumentation Grants S10RR029668 and S10RR027303. DMP thanks many collaborators and institutions for tissue samples processed in this study, and they will be recognized as co-authors or fully acknowledged in the resulting phylogenetic study in prep. DMP also thanks Sean Reilly and Ammon Corl for help in lab training and troubleshooting.

## Author Contributions

This experiment was designed by all authors, and DMP performed lab work with significant assistance from KB and LLS. KB developed a substantial proportion of bioinformatics pipelines, with contributions from DMP. DMP wrote the manuscript, with contributions from KB and LLS, and all authors approved the final manuscript.

## Data Accessibility

All the bioinformatics pipelines for transcriptome data processing and annotation are available at (https://github.com/CGRL-QB3-UCBerkeley/DenovoTranscriptome). Pipelines for marker development are available at (https://github.com/CGRL-QB3-UCBerkeley/MarkerDevelopmentPylogenomics). The bioinformatic pipelines of sequence capture data processing are available at (https://github.com/CGRL-QB3-UCBerkeley/denovoTargetCapturePhylogenomics). All custom python scripts for sequence capture performance evaluation are available on github (https://github.com/dportik/). These include tools for automating the calculation of coverage, duplication, sensitivity, specificity, and coverage uniformity. Additional scripts are available for evaluating and refining DNA sequence alignments. Molecular sequence data will be published in an associated manuscript in prep.

